# Beyond iron concentration: iron aggregation shapes quantitative MRI in the human brain

**DOI:** 10.64898/2026.05.19.726170

**Authors:** Alexander Stürz, Marlene Panzer, Bernhard Glodny, Elke R. Gizewski, Heinz Zoller, Christoph Birkl

**Affiliations:** Department of Radiology, Medical University of Innsbruck, Innsbruck, Austria; Neuroimaging Research Core Facility, Medical University of Innsbruck, Innsbruck, Austria; Department of Internal Medicine I, Medical University of Innsbruck, Innsbruck, Austria; Christian Doppler Laboratory for Iron and Phosphate Biology, Medical University of Innsbruck, Innsbruck, Austria

**Keywords:** Quantitative Susceptibility Mapping (QSM), Relaxometry, Quantitative MRI, Microstructure, Brain, Iron

## Abstract

Aceruloplasminemia (ACP) is a rare neurodegenerative disorder characterized by extreme cerebral iron overload and a shift towards larger iron aggregates, providing a unique possibility to study how iron aggregation shapes MRI contrast *in vivo*. We introduce a clinically feasible, multi-parametric quantitative MRI (qMRI) framework that combines quantitative susceptibility mapping (QSM), 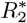, and *R*_2_ to disentangle changes in total iron concentration from alterations in iron aggregation and its spatial organization at the cellular scale. Our biophysical model links the microstructure sensitive 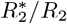 ratio and the slope of the susceptibility–relaxation relationship (*α*_iron_) to iron aggregation size and distribution. In a 3T qMRI study of three patients with ACP and three matched controls, we observe a marked increase in 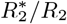 and a pronounced increase of the 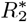–QSM slope (*α*_iron_: controls 154.09 *±* 52.89 s^−1^ppm^−1^; patients 296.68 *±* 57.18 s^−1^ppm^−1^; *p* = 0.016), consistent with enhanced iron aggregation and altered spatial organization. Model-based decomposition of transverse relaxation indicates that up to approximately 40% of the observed *R*_2_^∗^ elevation in ACP is attributable to changes in iron distribution beyond increased iron concentration alone. These findings establish a robust, translational qMRI approach for quantitative *in vivo* assessment of iron aggregation, revealing microstructural drivers of iron-related neurodegeneration that extend beyond bulk iron load.

## I. INTRODUCTION

Aceruloplasminemia (ACP) is a rare autosomal recessive neurodegenerative disease, representing one of the most severe forms of cerebral iron overload. It offers a unique opportunity to investigate the biological and biophysical consequences of pathological iron accumulation in the human brain *in vivo* [1, 2]. Loss-of-function mutations in the ceruloplasmin gene abolish the ferroxidase activity required to oxidize Fe^2+^ to Fe^3+^, thereby impairing transferrin-mediated iron export [3–5]. As a result, iron accumulates massively within glial cells, particularly astrocytes, most prominently in the basal ganglia, thalamus, and substantia nigra [4, 6]. Post-mortem iron quantification confirms markedly elevated total cerebral iron concentration in ACP [1, 7, 8]. Complementary histopathology shows that iron is not only increased in amount but is stored predominantly in highly aggregated intracellular forms, accompanied by cellular enlargement and astrocytic deformation [9–13].

Magnetic resonance imaging (MRI) enables noninvasive assessment of brain iron *in vivo* by exploiting its paramagnetic susceptibility, which induces local magnetic field inhomogeneities that alter both phase and relaxation of the MR signal [14, 15]. Among quantitative MRI (qMRI) techniques, the effective transverse relaxation rate *R*_2_^∗^, the irreversible transverse relaxation rate *R*_2_, and quantitative susceptibility mapping (QSM) are established and validated as iron-sensitive biomarkers [16–18].

QSM reconstructs bulk tissue magnetic susceptibility from gradient-echo phase using a macroscopic forward model [19, 20] and is often interpreted as a proxy for tissue iron content. In contrast, transverse relaxation is commonly described by empirically motivated effective rate constants and therefore provides a phenomenological, rather than fully mechanistic, description [21, 22]. Crucially, 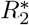 and *R*_2_ depend not only on the concentration of magnetic sources but also on their microstructural organization, such as size, clustering, and spatial distribution of para- and diamagnetic inclusions, and on diffusion mediated dephasing across multiple spatial scales [23, 24]. Physically grounded models that describe the impact of iron and other magnetic susceptibility sources on transverse relaxation are rare, as the governing mechanisms span multiple spatial scales dominated by distinct physical processes [24].

Most clinical qMRI studies to date have thus interpreted QSM, 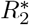 and *R*_2_ primarily as markers of tissue iron concentration, and have used them to map iron accumulation in neurological diseases including multiple sclerosis [25, 26], Parkinson’s disease [27, 28], and Alzheimer’s disease [29, 30]. However, treating *R*_2_^∗^ solely as a proxy for iron concentration is incomplete and overlooks its sensitivity to microstructural features that modulate transverse relaxation *in vivo* [22, 31]. In contrast, QSM predominantly reflects bulk susceptibility and is comparatively less sensitive to the microstructural configuration of underlying susceptibility sources. This complementarity suggests that combined susceptibility- and relaxation-based metrics can disentangle total iron concentration from alterations in iron aggregation and spatial organization [32].

Here, we investigate a clinically feasible, multiparametric qMRI framework in the context of extreme brain iron overload, aiming to separate iron load from aggregation-driven microstructural effects *in vivo*. Building on established approaches to characterize brain iron [32–35], we introduce two complementary microstructure-sensitive metrics: the 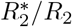 ratio and the slope of the 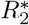-QSM relationship, *α*_Iron_. Our biophysical model links both metrics to iron aggregate size and spatial organization at the cellular scale. In ACP, these measures reveal substantially altered iron microstructure beyond increased iron content, and we show that microstructural effects account for up to approximately 40% of the observed increase in 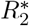 relative to matched controls (Fig. 1).

**FIG. 1.**
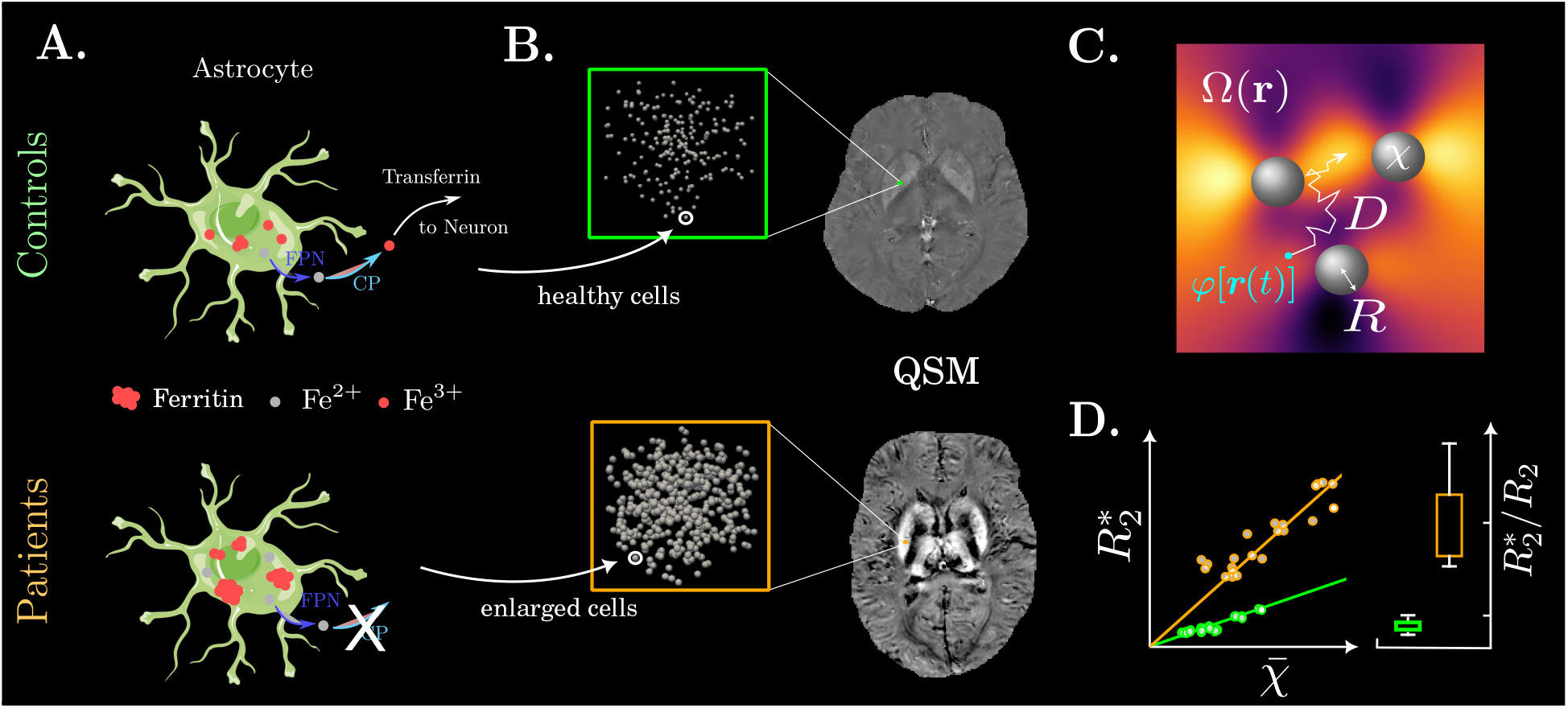
Conceptual overview of the proposed mechanism linking impaired iron handling to MRI contrast changes in aceruloplasminemia (ACP). **(A)** Under physiological conditions, ceruloplasmin (CP) enables iron export by oxidizing Fe^2+^ to Fe^3+^ for transferrin binding. In ACP, the absence of CP impairs iron export, leading to intracellular iron retention in cells, predominantly Astrocytes, where it is subsequently stored (mostly) in ferritin. **(B)** Macroscopic susceptibility maps (QSM) are shown for control and patient, alongside the corresponding changes in the underlying microstructural susceptibility distribution *χ*(***r***) in the zoomed-in regions. **(C)** Schematic of the model: protons accumulate phase *φ*[***r***(*t*)] while diffusing with coefficient *D* in the frequency field Ω(***r***) induced by spherical inclusions of radius *R* and intrinsic susceptibility *χ*. **(D)** Impact of microstructural alterations on macroscopic measurements of transverse relaxation and QSM.

Collectively, our results establish multi-parametric qMRI as a sensitive approach to assess microstructural iron pathology beyond total iron concentration. They further provide a physically interpretable framework for decomposing susceptibility–relaxation signatures into concentration versus microstructure, enabling more quantitative interpretation of iron-related MRI contrast in the context of iron-driven neurodegeneration [36].

## II. RESULTS

In patients with ACP, pronounced iron deposition is evident in 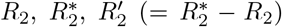 and QSM maps, most prominently within deep gray matter (Fig. 2). The largest changes occur in the putamen and, to a lesser extent, the thalamus, where signal increases are spatially heterogeneous. Cortical gray matter also shows marked signal increases, and white matter exhibits milder involvement, most apparent in *R*_2_^∗^ maps. The globus pallidus is least affected, with only subtle deviations from controls.

**FIG. 2.**
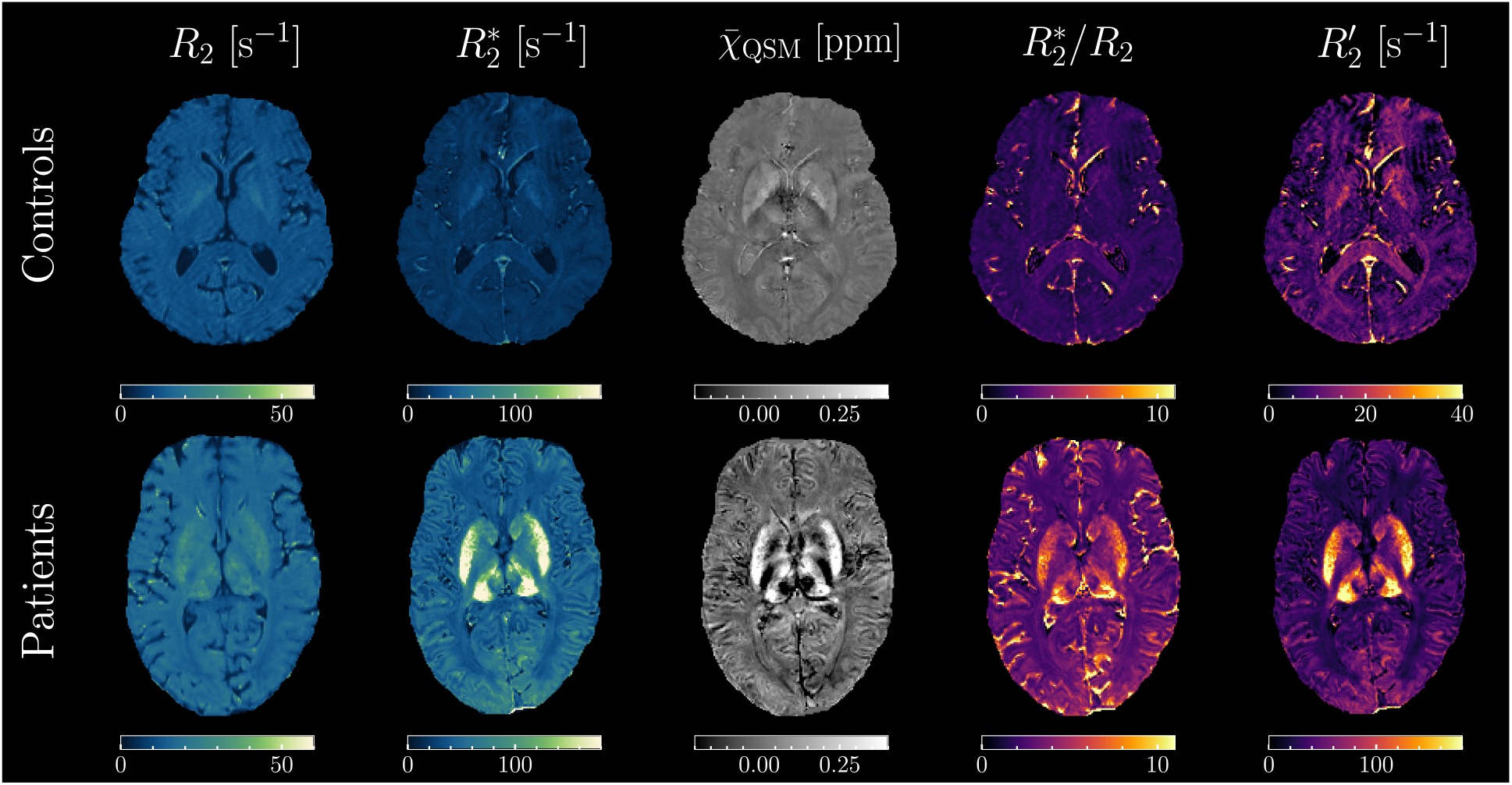
Quantitative *R*_2_, 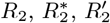, QSM and 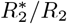 ratio maps of a healthy subject and a patient with aceruloplasminemia (ACP). All maps reveal pronounced iron overload in the deep gray matter and in the cortical gray matter of the patient compared to the control, while white matter appears largely unaffected. In the 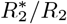 ratio map of the control subject, blood vessels dominate the contrast due to their strong static susceptibility effects. In the patient, contributions from the basal ganglia and thalamus—as well as smaller effects across the cortical gray matter—are also highlighted in addition to the vascular structures.

The 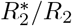 ratio is elevated in patients across deep gray matter compared to controls (Fig. 2). In healthy controls, high 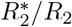 values correspond mainly to blood vessels, which are also hyperintense in 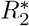 and QSM maps. In contrast, patients with ACP show elevated 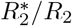 values throughout deep gray matter, indicating altered microstructure beyond vascular contributions.

ROI-based analysis revealed higher values of all qMRI parameters in patients compared to controls (Fig. 3, Table I). The largest increase in 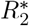 was observed in the putamen, with patient values of 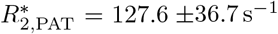, corresponding to an approximately ∼ 5.5-fold elevation over controls (*p <* 0.05). The smallest absolute 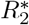 increase was observed in the globus pallidus, rising from 37.3 ± 4.1 s^−1^ in controls to 65.4 ± 10.4 s^−1^ in patients (*p <* 0.05). QSM showed a similar spatial pattern, with the largest increase in the putamen (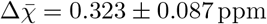 ppm; *p <* 0.05).

**Table 1.**
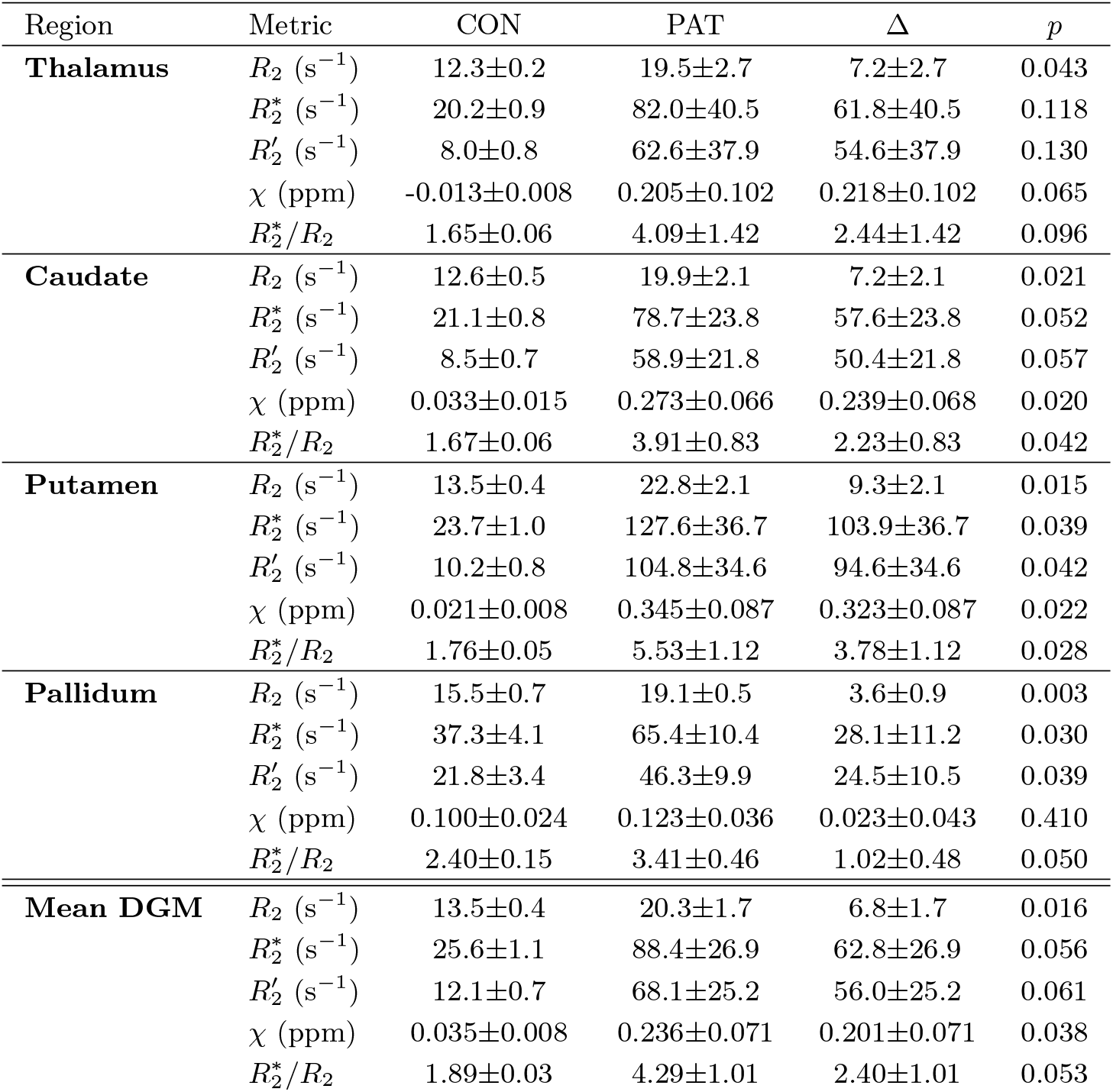
Regional relaxometric and susceptibility metrics (mean *±* SD) for controls (CON) and patients (PAT). Δ denotes the difference between patient and control.

| Region | Metric | CON | PAT | $\Delta$ | $p$ |
| --- | --- | --- | --- | --- | --- |
| **Thalamus** | $R_2$ (s<sup>-1</sup>) | 12.3±0.2 | 19.5±2.7 | 7.2±2.7 | 0.043 |
| | $R_2^*$ (s<sup>-1</sup>) | 20.2±0.9 | 82.0±40.5 | 61.8±40.5 | 0.118 |
| | $R_2'$ (s<sup>-1</sup>) | 8.0±0.8 | 62.6±37.9 | 54.6±37.9 | 0.130 |
| | $\chi$ (ppm) | -0.013±0.008 | 0.205±0.102 | 0.218±0.102 | 0.065 |
| | $R_2^*/R_2$ | 1.65±0.06 | 4.09±1.42 | 2.44±1.42 | 0.096 |
| **Caudate** | $R_2$ (s<sup>-1</sup>) | 12.6±0.5 | 19.9±2.1 | 7.2±2.1 | 0.021 |
| | $R_2^*$ (s<sup>-1</sup>) | 21.1±0.8 | 78.7±23.8 | 57.6±23.8 | 0.052 |
| | $R_2'$ (s<sup>-1</sup>) | 8.5±0.7 | 58.9±21.8 | 50.4±21.8 | 0.057 |
| | $\chi$ (ppm) | 0.033±0.015 | 0.273±0.066 | 0.239±0.068 | 0.020 |
| | $R_2^*/R_2$ | 1.67±0.06 | 3.91±0.83 | 2.23±0.83 | 0.042 |
| **Putamen** | $R_2$ (s<sup>-1</sup>) | 13.5±0.4 | 22.8±2.1 | 9.3±2.1 | 0.015 |
| | $R_2^*$ (s<sup>-1</sup>) | 23.7±1.0 | 127.6±36.7 | 103.9±36.7 | 0.039 |
| | $R_2'$ (s<sup>-1</sup>) | 10.2±0.8 | 104.8±34.6 | 94.6±34.6 | 0.042 |
| | $\chi$ (ppm) | 0.021±0.008 | 0.345±0.087 | 0.323±0.087 | 0.022 |
| | $R_2^*/R_2$ | 1.76±0.05 | 5.53±1.12 | 3.78±1.12 | 0.028 |
| **Pallidum** | $R_2$ (s<sup>-1</sup>) | 15.5±0.7 | 19.1±0.5 | 3.6±0.9 | 0.003 |
| | $R_2^*$ (s<sup>-1</sup>) | 37.3±4.1 | 65.4±10.4 | 28.1±11.2 | 0.030 |
| | $R_2'$ (s<sup>-1</sup>) | 21.8±3.4 | 46.3±9.9 | 24.5±10.5 | 0.039 |
| | $\chi$ (ppm) | 0.100±0.024 | 0.123±0.036 | 0.023±0.043 | 0.410 |
| | $R_2^*/R_2$ | 2.40±0.15 | 3.41±0.46 | 1.02±0.48 | 0.050 |
| **Mean DGM** | $R_2$ (s<sup>-1</sup>) | 13.5±0.4 | 20.3±1.7 | 6.8±1.7 | 0.016 |
| | $R_2^*$ (s<sup>-1</sup>) | 25.6±1.1 | 88.4±26.9 | 62.8±26.9 | 0.056 |
| | $R_2'$ (s<sup>-1</sup>) | 12.1±0.7 | 68.1±25.2 | 56.0±25.2 | 0.061 |
| | $\chi$ (ppm) | 0.035±0.008 | 0.236±0.071 | 0.201±0.071 | 0.038 |
| | $R_2^*/R_2$ | 1.89±0.03 | 4.29±1.01 | 2.40±1.01 | 0.053 |

**FIG. 3.**
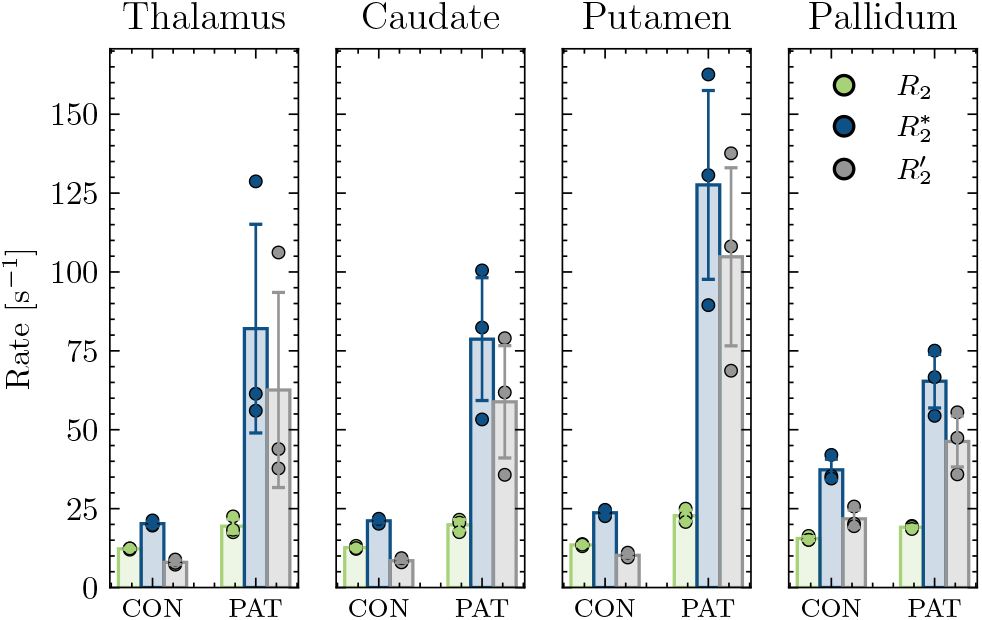
Group differences in transverse relaxation rates across the thalamus and basal ganglia. Barplots depict mean values of *R*_2_, 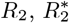, and the derived 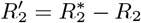, with individual dots representing subject-level measurements (three subjects per group). Error bars indicate the standard deviation across subjects.

*R*_2_ values were significantly elevated in patients in all analyzed brain regions (*p <* 0.05). The strongest increase in *R*_2_ was found in the putamen, rising from *R*_2,CON_ = 13.5 0.4 s^−1^ to *R*_2,PAT_ = 22.8 2.1 s^−1^ (*p <* 0.05). Overall, absolute patient-control differences were smaller for *R*_2_ than for 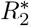 and QSM.

Consistent increases were also observed for 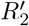 and for the 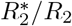 ratio across all assessed brain regions (Tab. I). The largest patient-control differences were again in the putamen, with an elevated 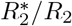 ratio of 5.53 ± 1.12 (*p <* 0.05) and a markedly higher reversible transverse relaxation rate 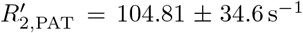 (*p <* 0.05).

### A. Elevated 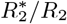 ratio indicates altered microstructure

To interpret these increases, we simulated microstructure-induced transverse relaxation as a function of magnetic inclusion size using Monte Carlo models of impermeable spherical inclusions for gradient-echo (Fig. 4 A) and spin-echo (Fig. 4 B) acquisitions (Sec. V H), guided by histological reports of aggregated intracellular iron in patients with ACP [1, 9]. We further computed 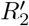 and the 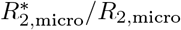 ratio (Fig. 4 C and D). Simulations were performed for different volume fractions of the magnetic inclusions, representing a changed total iron content, within a voxel.

**FIG. 4.**
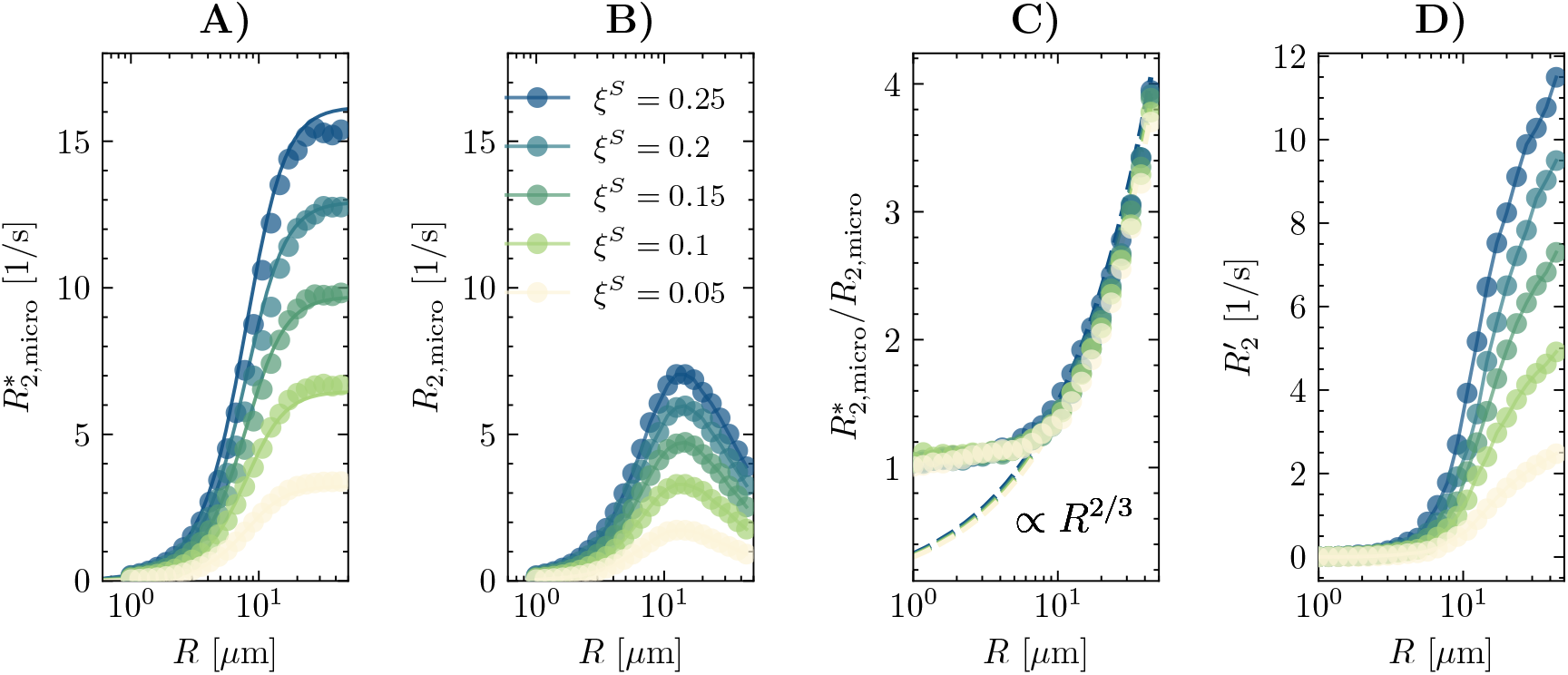
Simulated mGRE (**A**) and mSE (**B**) transverse relaxation rates for spherical magnetic inclusions with volume fraction *ξ*^*S*^ and intrinsic susceptibility *χ* = 4*π* · 10^−7^. The total volume fraction is distributed uniformly over spheres of radius *R*. (**C**) The ratio 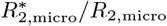 is invariant of volume fraction. (**D**) The reversible relaxation rate 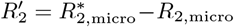 exhibits a clear dependence on the volume fraction of susceptibility inclusions. Simulations were performed using a diffusion coefficient of *D* = 1 *µ*m^2^*/*ms and a field strength of *B*_0_ = 3 T. The solid lines in (**A**) represent the Padé-approximation according to equation (12). All panels illustrate how increasing inclusion radius shift the relaxation behaviour toward static dephasing.

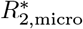 exhibited distinct asymptotic behavior at small and large inclusion radii approaching a plateau at larger radii. *R*_2,micro_ increased up to intermediate radii and decreased at larger radii, consistent with the static dephasing regime (SDR). Consequently, the 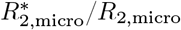 ratio increases systematically with inclusion radius and is independent of the magnetic inclusion volume fraction *ξ*^*S*^, in agreement with the analytic expression in equation (10), exhibiting the characteristic ∼ *R*^2*/*3^ scaling.

In contrast, *R*_2_^*′*^ depends on both inclusion size and volume fraction. The solid lines in Fig. 4 illustrate the Padé approximation given in equation (13), demonstrating its validity across the full range of simulated inclusion radii.

### B. Relaxation-Susceptibility Relationship reflects microstructurally compartmentalized iron

The relationship between 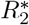 and QSM, averaged over all assessed deep gray matter regions, differed markedly between patients and controls (Fig. 5 B). We characterized this association via linear regression (Sec. V C), where the slope *α*_Iron_ reflects the susceptibility-dependent component of 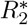∗, and the intercept represents the susceptibility-independent component, 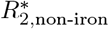 (equation (11)).

**FIG. 5.**
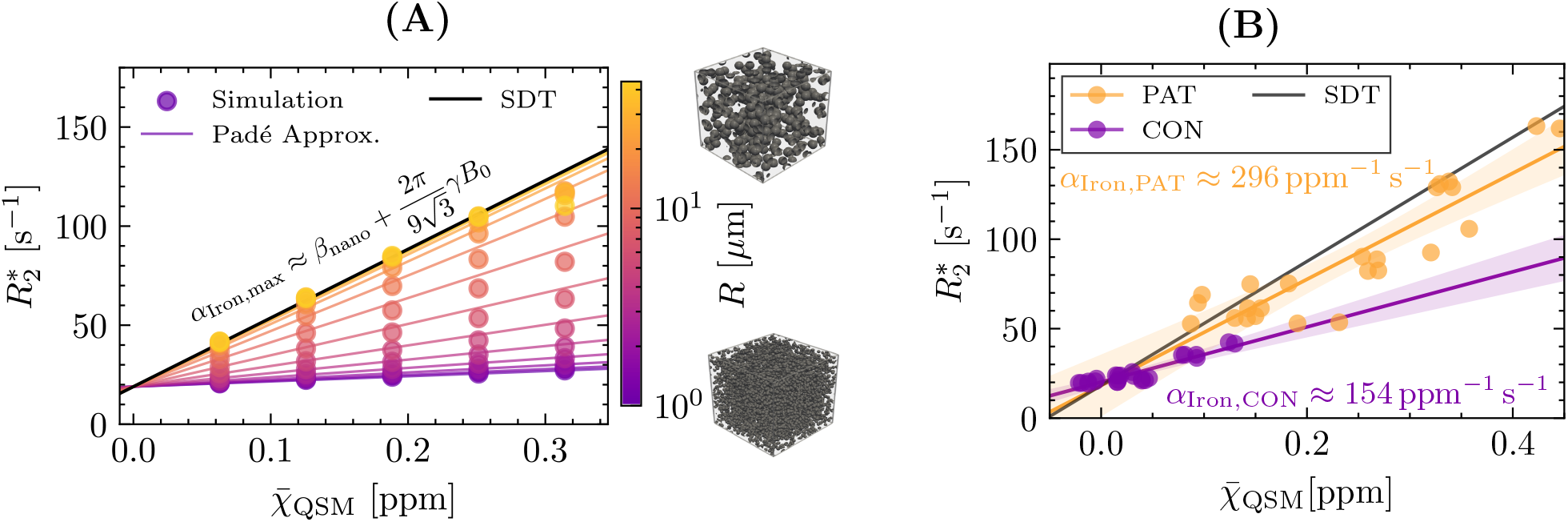
Relationship between 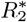 and QSM in deep gray matter. (**A**) Simulation of *α*_Iron_ for five different volume fractions *ξ*^*S*^ as a function of the radius *R*. Markers indicate the simulations, while the lines represent the theoretical model proportional to *α*_Iron_ = *β*_nano_ + *κ*(*R*), assuming 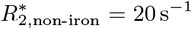. This highlights the dependence of *κ* on the inclusion radius, while intrinsic susceptibility and diffusion, which also influence *κ*, are kept constant. (**B**) In vivo measurements, where each point represents the mean value in the thalamus, caudate, putamen, and pallidum, separated by left and right hemisphere, for three patients (PAT) and three controls (CON). Orange markers denote patients, while violet markers indicate controls. The black curve represents the theoretical maximal slope predicted by static dephasing theory.

Simulations of the relaxation-susceptibility relationship (Fig. 5 A) illustrate the dependence of *α*_Iron_ on microstructural parameters, in particular the inclusion radius *R*.

The measured 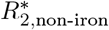 did not differ significantly between patients (18.25 ± 6.34 s^−1^) and controls (20.14 ± 2.94 s^−1^; *p* = 0.77). By contrast, *α*_Iron_ was substantially higher in patients (296.68 ± 57.18 s^−1^*/*ppm) than in controls (154.09 ± 52.89 s^−1^*/*ppm), yielding Δ*α*_Iron_ = 142.59 ± 57.18 s^−1^*/*ppm (*p* = 0.016). The joint regression model achieved an overall *R*^2^ = 0.930.

For comparison, the theoretical prediction from Static Dephasing Theory (black line in Fig. 5), gives *α*_Iron,SDR_ = 346.74 s^−1^*/*ppm, indicating that patient data lie considerably closer to the static-dephasing regime than control data.

### C. Decomposition of 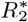 reveals strong dependence on microstructural iron organization

Using the biophysical model introduced in Sec.V D, measured 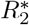 was decomposed into three components: non-iron related relaxation 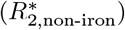, iron induced nanoscale relaxation 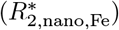, and iron induced microscale relaxation 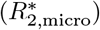. Averaged across all deep gray matter regions, patients and controls showed similar 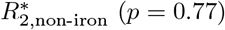, but different in both iron related components (Fig. 6). The average nanoscale iron contribution increased from 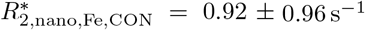 to 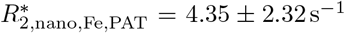 in patients (*p* = 0.04). The average microscale contribution showed a much larger difference, increasing from 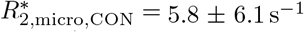 to 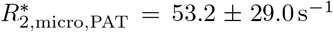, corresponding to a microstructure-to-nanoscale ratio of approximately 6.3 in controls and 12.2 in patients.

**FIG. 6.**
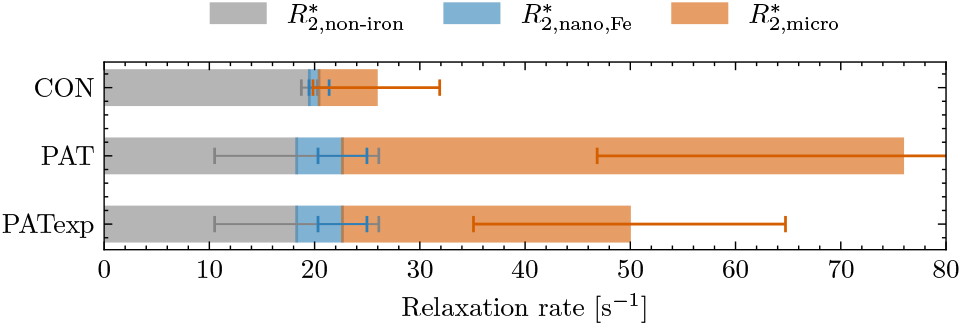
Decomposition of the average measured *R*_2_^∗^ across the segmented deep gray matter brain regions into: non-iron induced background 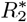, nanoscale iron-dependent relaxation 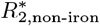, and microstructure-dependent relaxation 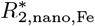. Here we present measured data from controls (CON) and patients (PAT), together with a theoretical patient condition assuming unchanged microstructure (PAT_exp_). PAT_exp_ represents the expected 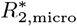 increase driven by iron-concentration changes alone, using *α*_Iron,CON_. The excess *R*_2_^∗^ observed in PAT beyond PAT_exp_ indicates an additional contribution from microstructure-dependent relaxation effects within the biophysical model.

To assess how much of the observed 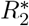 elevation could be explained by increased susceptibility (and thus iron concentration) alone, we computed a theoretical scenario (PAT_exp_) that preserves control microstructure while increasing susceptibility to patient levels, i.e. using *α*_Iron,CON_ to estimate 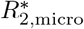. This yielded 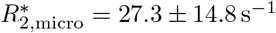.

Under these assumptions, approximately 62% of the measured increase in 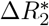 between patients and controls, is attributable to increased susceptibility alone, whereas the remaining 38% requires a change in microstructure in addition to increased iron content.

## III. DISCUSSION

In this study, we show that multi-parametric qMRI can disentangle iron microstructural effects from bulk tissue iron concentration *in vivo*. To achieve this, we introduce two complementary, microstructure-sensitive metrics: the 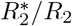 ratio and *α*_Iron_, the slope of the 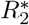–QSM relationship. Both metrics depend on the intrinsic susceptibility *χ*, the inclusion size *R*, and the diffusion coefficient *D*, while being independent of the volume fraction of susceptibility inclusions. In patients with ACP, both metrics indicate systematic alterations in the spatial organization of iron in addition to increased iron concentration. Our biophysical framework uses *α*_Iron_ to disentangle these two effects, thereby increasing the specificity of iron-related relaxation beyond conventional single-parameter approaches.

### A. 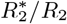 as a microstructure-sensitive mapping approach

The Monte Carlo simulations in Fig. 4 are consistent across a wide range of simulation frameworks, including magnetic susceptibility perturbations of various shape [35, 37–40]. Our simulations show that the 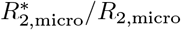 ratio is independent of the volume fraction of susceptibility inclusions over the full range of inclusion size *R* and follows the asymptotic behavior described by equation (10).

Experimentally, we measure total relaxation rates, 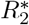 and *R*_2_, which include nanoscale relaxation components. Thus, the measured 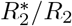 does not directly reflect the pure microscale ratio from the simulations in Fig. 4. However, as shown in Fig. 6, the nanoscale relaxation contribution differs by less, than 11% between controls and patients. Under these conditions, changes in the total 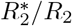 ratio predominantly reflect alterations in the microscale 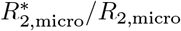 ratio.

Conceptually, this approach is similar to vessel size imaging [41], in that a ratio of rlaxation rates reports on microstructural length scales. Unlike vessel size imaging, however, the intrinsic susceptibility *χ* and the diffusion coefficient *D* cannot be assumed constant between patients and controls. Therefore, a direct, absolute quantification of the iron cluster size radius *R* is not yet feasible. Nontheless, as 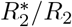 increases monotonically with *R* in the relevant regimes, group differences in the ratio provide robust evidence for underlying microstructural differences, including potential changes in iron cluster size and organization.

Together with the spatially coherent patterns in Fig. 2, the 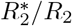 ratio offers a practical mapping contrast that is sensitive to microstructure and not solely to iron load. The observed distribution align with histopathological reports of intracellular, aggregated iron [1, 3, 9], underscoring the translational potential of this metric for neurodegenerative diseases characterized by abnormal iron aggregation.

### B. 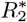 -QSM coupling as a marker of microstructure-dependent dephasing

The second metric, *α*_Iron_, is the slope of the relationship between bulk magnetic susceptibility and transverse relaxation. In our model (equation (11)), increases in *α*_Iron_ arise from either an increase in the microstructural coupling parameter *κ* or a change in the nanoscale relaxation component *β*_nano_.

Post-mortem studies indicate that ferrihydrite-bound iron is the predominant iron form in deep gray matter, accounting for approximately 91-95% of total iron in patients with ACP [7]. In healthy iron-rich deep gray matter, the majority of non-heme iron is likewise stored in ferritin (80–90%) [42], with spectroscopy-based estimates indicating at least 80% [43], while recent work reports values of up to 94% [44]. Given this dominant ferritin (ferrihydrite) pool in both groups, *β*_nano_ is expected to remain largely stable between patients and controls.

Minor fractions of other iron species, such as maghemite-bound iron, hemosiderin, and free Fe^3+^, have been detected by *Electron Paramagnetic Resonance* (EPR) and *Superconducting Quantum Interference Device* (SQUID) magnetometry in ACP brain tissue [7]. Fe^3+^ exhibits a magnetic moment of *µ* = 5.9*µ*_*B*_ [45], different from that of ferrihydrite. Moreover, histopathological studies qualitatively report elevated levels of neurotoxic Fe^2+^ (magnetic moment of *µ* = 4.9*µ*_*B*_), which cannot be stored in ferritin, [6, 12]. While in theory, these iron species could influence *β*_nano_ due to their different magnetic moments, their concentrations are several orders of magnitude lower than those of ferrihydrite-bound iron in both patients with ACP and controls [7, 8]. Any effect on *β*_nano_ is therefore expected to be negligible relative to the dominant ferritin (ferrihydrite) contribution. Within this context, group differences in *α*_Iron_ are most plausibly attributed to changes in the microstructural coupling parameter *κ*.

### C. Microstructural alterations affecting 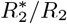 **and** *α*_**Iron**_

Our theoretical framework predicts that increases in both 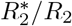 ratio and *α*_Iron_ can arise from three primary factors (equations (13) and (10)): larger effective inclusion size *R*, reduced diffusivity *D*, or increased intrinsic susceptibility *χ* of individual magnetic inclusions.

#### 1. Increase of Radius R

In the motional narrowing regime (MNR), the microstructural coupling parameter *κ* scales quadratically with the effective inclusion radius (equation (8)), making *κ* particularly sensitive to changes in radius compared to changes in intrinsic susceptibility or diffusion. Histopathological studies show hypertrophic iron-loaded astrocytes (up to 30–40*µ*m [1, 11]) and grumose foamy spheroid bodies (GFSBs) with diameters of 10– 60 *µ*m [9, 10] in the globus pallidus and putamen of patients with ACP. The size of these structures lies within the range known to induce static dephasing, similar to other microstructural contrast sources such as blood vessels (capillary radii of 2.5–25 *µ*m [41]), dopaminergic neurons in the substantia nigra (∼14 *µ*m) [34, 35], and spherical polystyrene microbeads (20–40 *µ*m) in phantoms [40].

#### 2. Increase of intrinsic susceptibility χ

Intrinsic susceptibility, reflecting the magnetic properties of individual inclusions, may increase with higher iron loading due to impaired cellular iron efflux (see Fig. 1). The substantial increase in iron concentration observed in various post-mortem studies is likely to accumulate intracellular, thereby promoting iron compartmentalization and consequently enhancing intrinsic susceptibility [1, 7, 8]. A similar mechanism has been demonstrated *in vitro* in (U)SPIO-loaded THP-1 macrophages, where strong iron compartmentalization resulted in static dephasing due to an increase in intrinsic susceptibility [46]. However, these alterations in cellular iron content in ACP are accompanied by pronounced cellular hypertrophy and enlargement of iron-containing structures [9]. As iron accumulation becomes distributed over an increasing radius, a higher total cellular iron load does not necessarily lead to a proportional increase in intrinsic susceptibility.

#### 3. Reduced diffusivity D

Cellular hypertrophy and increased packing density (e.g. within GFSBs) suggest progressive structural compaction and the formation of diffusion barriers [1, 11], which would reduce *D* and increase *κ*. Although diffusion metrics were not assessed in the present study, typical mean diffusivity (MD) in deep gray matter is in the range of *D* = 0.7 − 1.0 *µ*m^2^*/*s [32, 47] and can decrease in pathologies associated with elevated cellular density, such as gliomas[48]. Whether the pronounced susceptibility-induced dephasing observed in ACP would permit robust diffusion measurements remains an open question, as the strong microstructure-induced field gradients may substantially interfere with the effects of the external diffusion gradients and thereby bias the apparent diffusion metrics [49, 50].

The observed increase in *α*_Iron_, mediated through *κ*, as well as the increase in the 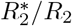 ratio, is most consistently explained by a combination of all three mechanisms. We propose, that changes in the effective inclusion radius *R* represent the dominant factor, based on (i) the strong dependence of *κ* on *R* (equation (13)), (ii) the sensitivity of 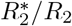 to changes in the effective inclusion radius *R* (equation (10)), and (iii) concordant histopathological evidence of enlarged iron-containing structures [1, 9, 51].

Our approach differs from Yablonskiy et al. [33], who explicitly separate heme and non-heme iron, by instead, modeling the aggregate effect of all microscopically distributed susceptibility sources on transverse relaxation. Conceptually, it aligns more closely with the iron microstructural coefficient introduced by Taege et al. [32]. To our knowledge, this is the first *in vivo* evidence that microstructural alterations drive the 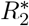-QSM relationship (*α*_Iron_) towards its theoretical limit in deep gray matter of patients with ACP.

### D. Limitations

While our biophysical model ensures computational robustness, several limitations warrant consideration. First, the 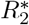∗–QSM slope can be estimated reliably, however, the corresponding intercept is sensitive to the specific tissue regions selected for analysis.

On the technical side, isotropic whole-brain mapping of *R*_2_ remains challenging due to the lack of accelerated 3D multi-echo spin echo sequences on most clinical MRI systems. We therefore computed *R*_2_ using two echoes, which is a common approach, although it may lead to an underestimation of *R*_2_ values [24, 52].

Furthermore, a limitation arises from the macroscopic nature of QSM, which assumes a negligible microstructure-induced Larmor frequency shift 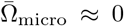 (often referred to as the mesoscopic Larmor frequency shift). While this assumption is valid for spherical inclusions in the MNR, a dependence 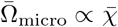 may emerge in the SDR for *t* ≫ *τ*_*D*_ [53, 54].

Finally, our model does not explicitly account for heme iron contributions, which may serve as a confounding factor in susceptibility-based measurements [33]. However, given the underlying biology of ACP, characterized by intracellular iron retention rather than hemolysis [5], a dominant contribution of heme iron to the observed differences between patients and controls is unlikely, but cannot be fully excluded.

Additionally, rapid signal decay and the choice of susceptibility reference have posed significant challenges for qMRI in post-mortem studies [8]. While recent *in vivo* work has demonstrated the feasibility of QSM in ACP [55], susceptibility referencing remains a critical source of variability, potentially affecting the interpretation of relaxation metrics beyond iron-specific contributions.

## IV. CONCLUSION

By linking magnetic susceptibility and transverse relaxation through a biophysical framework, we demonstrate that multi-parametric qMRI can separate microstructural alterations in iron organization (e.g., aggregate size and spatial arrangement) from changes in total iron concentration. Applied *in vivo* in a setting of extreme iron accumulation, the framework yields findings that are consistent with histopathology and substantially enhance the interpretability of iron-sensitive MRI.

Quantitatively, we show that in patients with ACP, up to ∼ 40% of the observed increase in 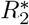 can be attributed to microstructural effects beyond increased iron concentration alone. This implies that conventional 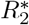 -based assessments may overestimate iron content if microstructural effects are not considered. The proposed approach, based on clinically feasible sequences and straightforward modeling, can be easily integrated into existing MRI protocols. Because the model addresses generic mechanisms of susceptibility-driven dephasing, it is readily extensible to other conditions with iron accumulation or altered susceptibility sources. These microstructure-sensitive relaxometric measures offer a path to investigate, monitor, and ultimately target the biological mechanisms of iron-related neurodegeneration *in vivo*.

## V. METHODS

For this study, three female controls (aged 30–51 years) and three female patients diagnosed with ACP (aged 50–52 years) were examined.

### A. Data acquisition

MRI was performed on a 3 T system (MAGNETOM Skyra, Siemens Healthineers, Erlangen, Germany) using a 64-channel head-and-neck receive coil. The imaging protocol included following sequences:

- a magnetization prepared rapid gradient echo (MPRAGE) sequence for anatomical overwiew and tissue segmentation (*T*_*E*_ = 2.12 ms, *T*_*R*_ = 1690 ms, inversion time *T*_*I*_ = 900 ms, flip angle = 8^°^, isotropic resolution = 0.8 mm^3^).
- a 3D multi-echo gradient-echo (mGRE) sequence for 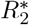 mapping and QSM (9 echoes, *T*_*E*_ = 2.6*/*5.2*/*7.8*/*10.4*/*13.0*/*15.6*/*18.2*/*20.8*/*23.4 ms, *T*_*R*_ = 36 ms, bandwith = 455 Hz*/*pixel. flip angle *α* = 15^°^, isotropic resolution = 1 ×1 ×1 mm^3^).
- a 3D turbo spin-echo sequence for *R*_2_ mapping (two acquisitions with effective echo times = 13 ms and 122 ms, repetition time *T*_*R*_ = 1000 ms, isotropic resolution = 1 × 1 × 1 mm^3^).

### B. Image processing

*T*_1_-weighted images were registered to the mGRE space using FSL FLIRT, and subcortical structures (Putamen, Pallidum, Caudate, Thalamus) were segmented with FSL FIRST [56]. For 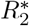 calculation we incorporated macroscopic field correction [57] and Rician noise-floor correction [34]. QSM reconstruction was performed according to the consensus guidelines [58], included ROMEO phase unwrapping [59], V-SHARP background field removal [60], and iLSQR dipole inversion [60], with susceptibility values referenced to the CSF. *R*_2_ values were obtained using a linear fit in log-signal space.

### C. Statistical analysis

For ROI-based comparisons of qMRI metrics between patients and controls, Welch’s t-test was applied to account for potential unequal variances between groups.

Given the small sample size (*n* = 3 per group), normality assumptions cannot be reliably evaluated and statistical significance should be interpreted in a descriptive manner.

For the analysis of the relationship between 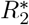 and 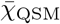, region-of-interest (ROI) mean values were extracted separately for left and right hemispheric structures (*N* = 24 data points per group). Because multiple ROIs originate from the same subjects, these observations are not strictly statistically independent. Nevertheless, to characterize the susceptibility–relaxation coupling within this mechanistic framework, a multiple linear regression model with interaction was fitted:

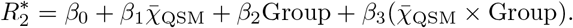

In this formulation, *β*_0_ represents the baseline relaxation offset 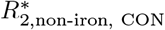, *β*_1_ the susceptibility–relaxation coupling coefficient in controls *α*_Iron,CON_, *β*_2_ a group-dependent shift in baseline relaxation 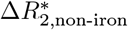, and *β*_3_ the group-dependent change in susceptibility–relaxation coupling between patients and controls Δ*α*_Iron_.

### D. Biophysical Model

The biophysical model employed in this work can be derived by considering the MR signal within a voxel of volume *V*, located at position ***R*** and observed at time point *t*. The signal is defined as the ensemble average over all spins within the voxel, each characterized by a phase *φ*.

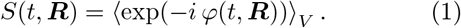

Since the underlying phase distribution is generally unknown, the signal can be expressed using a cumulant expansion. To leading order, this yields two dominant contributions: the mean phase ⟨*φ*⟩, which constitutes the primary observable in QSM, and the phase variance ⟨*φ*^2^⟩ − ⟨*φ*⟩^2^, which governs transverse relaxation and thus forms the main observable in relaxometry. Higher order cumulants can not be neglected in general.

#### 1. Quantitative Susceptibility Mapping

QSM infers the underlying bulk magnetic susceptibility distribution from the MRI mean-phase signal. The measured mean phase ⟨*φ*⟩, typically sampled across multiple echo times, reflects the voxel averaged Larmor frequency 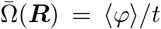. This frequency shift is assumed to arise from long-range magnetic field perturbations generated by a spatially varying macroscopic bulk susceptibility 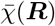(***R***) relative to a reference region [19, 20]. Micro- or mesoscopic contributions of Larmor frequency shifts are neglected 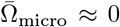. These macroscopic field inhomogeneities can be modeled by a convolution of the susceptibility distribution with the magnetic point dipole ϒ(***R***),

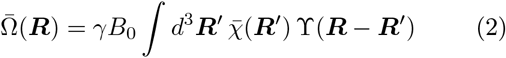

which describes the long-range contribution of susceptibility sources to the observed phase.

For deep gray matter structures, where most of the non-heme iron is stored in ferritin [44], the macroscopic model leads to the linear relationship of the bulk suscpetibility and the tissue iron concentration.

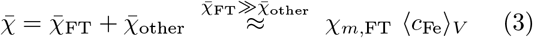

Here, *χ*_*m*,FT_ denotes the mass susceptibility of ferritin, and ⟨*c*_Fe_⟩_*V*_ the voxel-averaged iron concentration.

#### 2. Relaxometry

Transverse relaxation is reflected as the phase-variance in the ensemble average of equation (1), characterized by the effective transverse relaxation rate 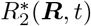. The observed relaxation rate comprises contributions arising from different spatial length scales [24] and is in general time-dependent.

For GRE acquisitions, the total relaxation rate can be decomposed as

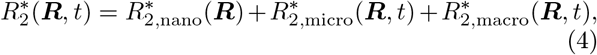

where the nanoscale term reflects molecular effects, the microscale term captures dephasing induced by mesoscopic magnetic structures in the presence of diffusion, and the macroscopic term accounts for large-scale field inhomogeneities.

#### 3. Nanoscale Relaxation

The nanoscale relaxation is based on short-range interactions. We can distinguish between two major contributions: iron-induced nanososcale relaxation, that we call 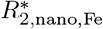, dependent on the iron binding site [61] and non-iron induced nanoscale relaxation 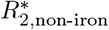, like spin-spin interactions [34]. The nanoscale iron relaxation is directly connected to the iron concentration via 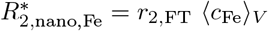 with *r*_2,FT_ being the relxivity of ferritin.

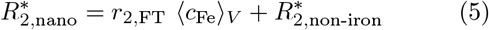

For *r*_2,FT_ = 0.0099 s^−1^*/*(*µ*g*/*g) we used the phantom evaluated relaxivity of ferritin at 7T and interpolated to our field strength of 3T [62]. This allows a conncetion between bulk and nansocale relaxation with the value *β*_nano_ = *r*_2,FT_*/χ*_*m*,FT_ ≈ 22.34 s^−1^*/*ppm. The mass susceptibility of ferritin *χ*_m,FT_ is described in Sec. V G.

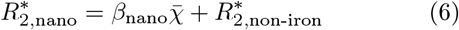

Transverse relaxation induced by magnetic tissue microstructure have been the subject of extensive investigation (see Kiselev and Novikov [24]). A central problem is understanding how diffusion impacts the accumulation of phase in the perturbed magnetic field generated by microscopic susceptibility variations *χ*(***r***). Note that ***r*** denotes the sub-voxel position on the microscale, embedded in a certain MR-visible medium (mostly water).

Transverse relaxation is then given by the sum of al real cumulants described in equation (1), corresponding to the phases accumulated along a diffusion path ***r***(*t*) in the frequency field Ω(***r***).

Obtaining analytic formulations for transverse relaxation induced by spatial susceptibility distributions at the microstructural level is generally challenging and is only possible in two limiting regimes, namely the SDR and MNR. In the SDR, where diffusion effects are negligible compared to the variance of Larmor frequency perturbations, such that *δ*Ω · *τ*_*D*_ ≫ 1, where *δ*Ω denotes the *standard deviation* of Larmor frequency variations and *τ*_*D*_ the characteristic diffusion time. In this regime, the measured ensemble average reflects the Fourier transform of the distribution of Larmor frequency shifts, such that the signal is directly governed by the underlying frequency distribution. Consequently, higher-order cumulants in equation (1) contribute significantly to the signal behavior. In this regime, a direct relationship between bulk magnetic susceptibility, as measured by QSM, and transverse relaxation can be established, as described by static dephasing theory (equation (7)) [54].

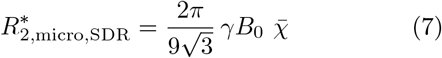

In the opposite limiting case, diffusion dominates over Larmor frequency perturbations, such that *δ*Ω · *τ*_*D*_ ≪ 1. This regime is commonly referred to as the MNR. For impermeable spherical inclusions, the relationship between transverse relaxation and magnetic susceptibility is given by equation (8) [63, 64], where the characteristic diffusion time is defined as *τ*_*D*_ = *R*^2^*/*(6*D*). The exact form of this relationship depends on the geometry of the inclusions as well as their permeability. In this study, impermeable spherical inclusions were assumed to provide a good approximation.

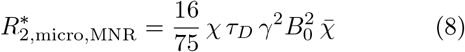

In the motional narrowing regime, the relationship between bulk magnetic susceptibility and transverse relaxation remains linear. As demonstrated by the simulations in Fig. 4 and Fig. 5 A, the microstructural contribution satisfies 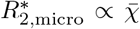, and is independent of the volume fraction of magnetic inclusions. However, it depends explicitly on microstructural parameters, including the intrinsic susceptibility of the inclusions *χ*, their radius *R*, and the diffusion coefficient *D*. Consequently, the linear coupling between 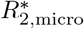 and 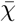 is characterized by the coupling constant *κ*. Dependent on the regime, the coefficients vary *κ*_MNR_ and *κ*_SDR_.

#### 5. Spin-Echo Relaxation

The same length-scale decomposition as in equation (4) applies to the spin-echo relaxation *R*_2_. Due to the refocusing pulse, macroscopic dephasing is suppressed and differences to 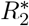∗ arise from microstructural effects (see Fig. 4). In the slow-diffusion regime, spinecho relaxation follows the scaling in equation (9) [41].

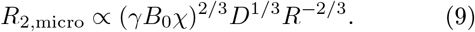

### E. Multi-parametric qMRI

Multi-parametric qMRI analysis was divided into two parts. First, the relationship between 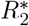 and *R*_2_ was examined. Here, *R*_2_ denotes the spin-echo relaxation rate and, similar to equation (4), consists of nanoscale and microscale relaxation components with 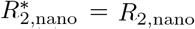. Combining equation (7) and equation (9), we find

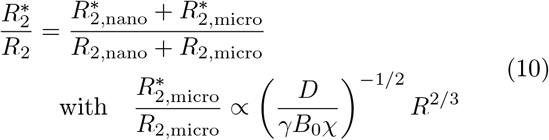

The behaviour in equation (10) is independent on the volume fraction of magentic inclusons *ξ*^*S*^ and is sensitive to microstructural parameters of diffusion *D*, intinsic susceptibility *χ* and the radius *R*. An alternative combination is given by the commonly used 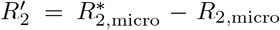, which is independent of the nanoscale relaxation contribution, but does not eliminate the influence of *ξ*^*S*^. Fig. 4 C shows the ratio of microstructure-induced relaxation rates validating equation (10).

Second, motivated by the concept of an iron microstructural coefficient [32], we express the total effective transverse relaxation rate as a combination of equations (4), (12), and (6).

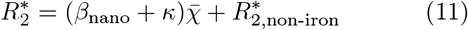

Here, *κ* represents the susceptibility-dependent relaxation coefficient defined in equation (13). We therefore define the total iron-related slope as *α*_Iron_ = *β*_nano_ + *κ*, comprising contributions from both nanoscale and microscale iron-induced relaxation mechanisms. Notably, the maximum slope at 3 T is obtained in the SDR, yielding *α*_Iron_ = 346.74 s^−1^*/*ppm.

### F. Dependency of *κ* on microstructural parameters

To obtain a formulation of *κ* across the full dynamic range, interpolating between MNR and SDR, we employ a Padé approximation,

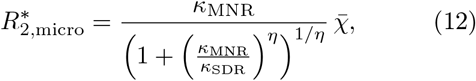

where *η* controls the smoothness of the transition between regimes. Rearranging yields an explicit expression for *κ*,

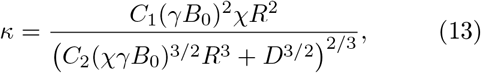

We chose *η* = 3*/*2, as this value provided the best empirical interpolation for our simulations (see Fig. 4), where we simulated 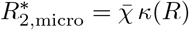. The constants were determined as *C*_1_ ≃ 0.0296 and *C*_2_ ≃ 0.0199.

### G. Mass susceptibility of Ferritin

The frequently cited magnetic moment per Fe atom in a ferrihydrite crystal, 3.8 *µ*_B_ [15], has been subject to debate in recent years [65, 66]. This value was derived under the assumption that each Fe atom in a ferrihydrite crystal contributes an identical magnetic moment that can be added linearly according to Curie’s law. Such an approach has been criticized for oversimplifying the magnetic behavior of ferritin and neglecting its complex internal structure. In particular, the magnetic susceptibility of ferritin is not governed by a single paramagnetic contribution, but instead arises from a superposition of different magnetic responses. These include a superparamagnetic contribution originating from uncompensated antiferromagnetically coupled spins within the ferritin core *M* ^SPM^, as well as a Curie–Weiss–type paramagnetic contribution associated with weakly coupled Fe atoms *M* ^PM^, primarily located at the surface [67].

In this work, we adopted the model proposed by Brooks [67] *M* = *M* ^SPM^ + *M* ^PM^ and obtained a mass susceptibility of *χ*_*m*,FT_ ≈ 0.443 ppb*/*(*µ*g*/*g) for a given iron concentration *c*_Fe_, according to equation (14).

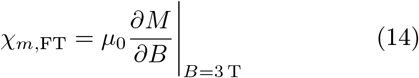

### H. Monte Carlo Simulation

To investigate the effects of micrstructural parameters on spherical perturbers, we performed Monte Carlo simulations of spherical magnetic inclusions [34, 38, 39, 68, 69] which serves to validate the theoretical hypotheses of equation (13) and to assess the influence of microstructure-induced spin-echo and gradient-echo relaxation rates.

The simulation was implemented on an *n* × *n* × *n* grid with *n* = 700. We placed randomly distributed spherical objects of radius *R*, occupying a volume fraction *ξ*^*S*^. The Larmor frequency shift Ω(***r***) was computed from the susceptibility distribution using a dipole convolution Ω(***r***) = *γB*_0_ℱ^−1^ {ϒ(***k***)*χ*(***k***)}.

Proton motion was modeled using Gaussian diffusion, 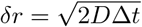, with multiple substeps per time interval. Boundary conditions were periodic on the domain, and rejection sampling was used to model collisions with impermeable spheres.

Signal formation inside the spheres was not simulated. The MR signal was computed from the accumulated phase along Gaussian diffusion paths according to equation (1). The gradient-echo relaxation rate was evaluated at *t* = 60 ms, and the spin-echo relaxation rate at *t* = 100 ms. For the spin-echo simulation, a *π*-pulse (spin flip) was applied at *T*_*E*_*/*2. Linear sub-grid interpolation is used. The simulation was checked for convergence and validated against the analytical limits given in equations (8), (7). The implementation was written in C++ with OpenMP parallelization [70], using 1×10^6^ simulated protons.

## ACKNOWLEDGMENTS

We thank Melanie Bauer and Christian Kremser for valuable discussions and helpful insights that supported the development and interpretation of this work. This research was funded in whole, or in part, by the Austrian Science Fund (FWF) [10.55776/I6838]. Images were provided by Servier Medical Art (https://smart.servier.com), licensed under CC BY 4.0.

## VI. AUTHOR CONTRIBUTIONS

**Alexander Stürz:** Conceptualization, Methodology, Software, Formal analysis, Investigation, Data curation, Visualization, Writing – original draft. **Marlene Panzer:** Investigation, Data curation, Resources, Writing – review & editing. **Bernhard Glodny:** Resources, Investigation, Writing – review & editing. **Heinz Zoller:** Investigation, Resources, Validation, Writing – review & editing. **Christoph Birkl:** Conceptualization, Methodology, Supervision, Project administration, Writing – review & editing. **Elke R. Gizewski:** Resources, Investigation, Writing – review & editing.

## VIII. DATA AVAILABILITY STATEMENT

The imaging data supporting the findings of this study contain sensitive patient information and are therefore not publicly available. Anonymized data may be made available from the corresponding author upon reasonable request and subject to institutional and ethical approval.

## IX. CODE AVAILABILITY STATEMENT

The custom Monte Carlo simulation code used to generate the results reported in this study is available at https://github.com/q373as/MonteCarloSimulationSpheres (version v1.0).

